# More than Drylands but Less of Dry Forests

**DOI:** 10.1101/147942

**Authors:** Marcelino de la Cruz, Pedro F. Quintana-Ascencio, Luis Cayuela, Carlos I. Espinosa, Adrián Escudero

## Abstract

The recent Global Drylands Assessment of forests is based on both an incomplete delimitation of dry forests distribution and on an old and incorrect delimitation of drylands. Its sampling design includes a large proportion of plots located in humid ecosystems and ignores critical areas for the conservation of dry forests. Therefore its results and conclusions are unreliable.

## Main Text

The recent Global Drylands Assessment of forests (*1*) clearly responds to a worldwide concern for better management and conservation of these ecosystems. Uncertainties about the contribution of these vast ecosystems covering more than 40 % of Earth’s surface to the global carbon balance and the value of their services make these estimations necessary and timing. The presumed significant underestimates of forest cover presented by Bastin et al. (*1*) have enormous implications for strategies of mitigation and adaptation to global warming. However, a carefully analysis of their data and methodology reveals significant drawbacks limiting the reach of these conclusions and their implications.

First, they claim that they use “the delineation adopted by the United Nations Environment Programme World Conservation Monitoring Centre [(UNEP-WCMC)]—lands having an aridity index (AI) lower than 0.65” (*2*). This affirmation is wrong. The delineation of UNEP-WCMC was based both in the United Nations Convention to Combat Desertification (UNCCD), i.e., the AI criterion, plus on the addition of some ecoregions (*3*) or “relevant areas” from the point of view of the Convention on Biological Diversity(CBD). As Bastin et al. (*1*) map in Fig. S1 shows, they ignored (areas in dark grey) both zones “presumed included” and zones “to review” in the UNEP-WCMC proposal. Therefore, they are using UNCCD criterion (i.e., AI <0.65), and not UNEP-WCMC delineation. This represents a step back in studying the relationship between desertification and biodiversity loss.

This choice has had two serious consequences. First, that the “global dryland extent” considered was based on UNEP-GRID (*4, 5*) aridity zones dataset. This is an old climatic database, with some errors, that induced Bastin et al. (*1*) to sample within non-dry areas. Even if they would have desired to delineate drylands based exclusively on AI, they should have used a corrected version (*6*) or, even better, they should have computed it by themselves. Computing AI based on the standard global database used extensively by ecologists and biogeographers (i.e., worldclim, *7-9*) shows that a significant proportion of the plots evaluated by Bastin et al. (*1*), around 22000 plots (i.e., a 10 %), were instead placed in areas with AI ≥ 0.65, i.e., non-dry areas according to UNCCD. These mistakes become apparent with a superficial inspection of the Global Drylands Assessment. For instance, most of the Ecuadorian easternmost rainforests (e.g. the Yasuní forest in the Napo region) and extensive parts of the south-eastern Peruvian Amazonian forests were included as drylands. This error is also apparent comparing the UNCCD aridity map (*2*) and is corrected version (*6*). This is a serious issue for the assessment of global dry forest cover because, for example, there are significant differences in tree cover between plots in non-dry (average 0.47) and genuine dry (average 0.11) locations (Kruskal-Wallis χ^2^ = 15233, df = 1, p-value < 0.0001).

There is a second and not least serious caveat, consequence of the drylands delimitation employed, which is the exclusion of the “relevant areas” for CBD (*2*), including the seasonal dry forests. In fact, among the 867 WWF terrestrial ecoregions (*3*), 54 (i.e., around a 5%) represent different aspects of the “Tropical and Subtropical Dry Broadleaf Forests” biome. This exclusion is evident when one overlays the map of ecoregions (*3*) with the map of dry areas employed by Bastin et al. (*1*). At least 52% of the area considered by WWF as dry forest is not contained in UNCCD delineation of drylands (*2*) and therefore it is not considered by Bastin et al. (*1*). As a consequence, some critical dry regions were omitted (*10, 11*). A simple comparison between them shows, for instance, the exclusion of extensive regions in Central America and Mexico (Fig. 1) and from most of the Asian South West. On the contrary, the strict application of the erroneous AI map without further considerations, resulted in some awkward facts such as including 56% of world flooded grasslands, 10% of moist broadleaf forests, 23% of mangroves and 51% of temperate forests as “dryland areas”. In fact, placing the studied plots over the ecoregions map (*3*) shows that almost 35000 plots (i.e. around 17%) fall within “non-dryland” biomes, including 11887 plots located in moist forests (5.6 %), 13313 in temperate forests (6.2 %), 5192 in taiga forests (2.4 %), and even 362 plots in mangroves (Table 1). All this makes Bastin et al. (*1*) study irrelevant for the assessment of dry tropical forest extent.

**Fig. 1.**
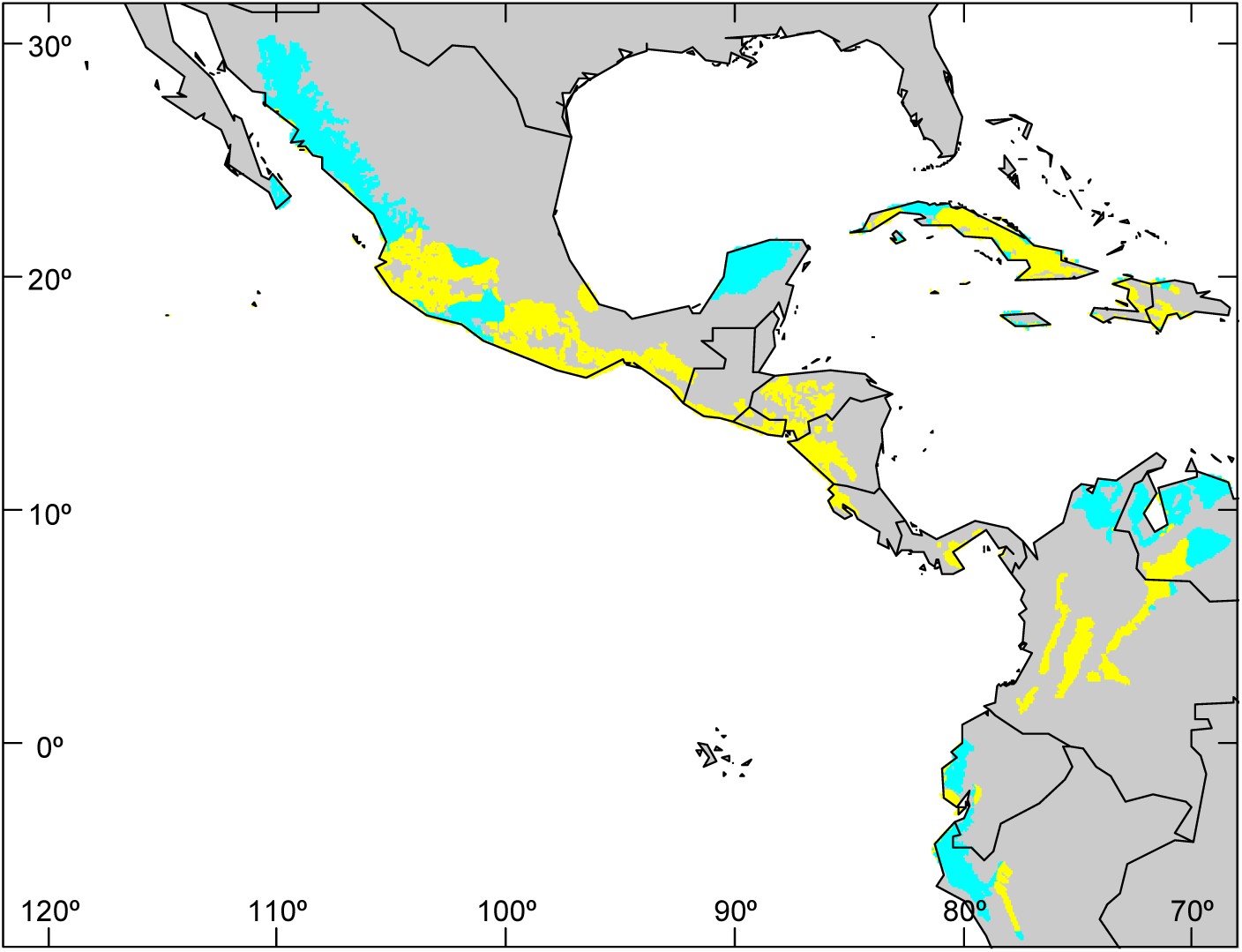
Distribution of Tropical and Subtropical Dry Broadleaf seasonal Forests in North, Central and part of South America (*3*). Cyan: included in UNEP-WCMC (*2*) and Bastin et al. (*1*) definition of drylands. Yellow: Not included.

**Table 1.** Distribution of the 213795 sampling plots of Bastin et al. (*1*) over WWF terrestrial biomes (*3*) and over a global map of aridity index (AI) computed on modern climatic databases (*7-9*). Figures in each cell represent number of plots within each combination (biome x AI category). Two plots were not considered because of incomplete coordinates in the database of Bastin et al. and another 142 had no available data in aridity index map.

|  | % plots /biome | Aridity index (AI) |  |
| --- | --- | --- | --- |
|  |  | AI ≥0.65 | AI <0.65 |
| NON DRY BIOMES |  |  |  |
| Tropical and Subtropical Moist Broadleaf Forests | 5.56 | 6710 | 5177 |
| Tropical and Subtropical Coniferous Forests | 0.33 | 137 | 578 |
| Temperate Broadleaf and Mixed Forests | 4.68 | 3278 | 6717 |
| Temperate Conifer Forests | 1.55 | 1112 | 2206 |
| Boreal Forests/Taiga | 2.43 | 3529 | 1663 |
| Flooded Grasslands and Savannas | 1.42 | 315 | 2718 |
| Tundra | 0.12 | 254 | 2 |
| Mangroves | 0.17 | 202 | 160 |
| Class 98 (inland water) | 0.04 | 17 | 74 |
| Class 99 (rock and ice) | 0.01 | 27 | 0 |
| TOTAL | 16.31 | 15581 | 19295 |
| **DRY BIOMES** |  |  |  |
| Tropical and Subtropical Dry Broadleaf<br>Forests | 3.25 | 1297 | 5649 |
| Tropical and Subtropical Grasslands,<br>Savannas and Shrublands | 25.45 | 2175 | 52197 |
| Temperate Grasslands, Savannas and<br>Shrublands | 11.62 | 1152 | 23676 |
| Montane Grasslands and Shrublands | 2.59 | 508 | 5019 |
| Mediterranean Forests, Woodlands and<br>Scrub | 4.66 | 306 | 9644 |
| Deserts and Xeric Shrublands | 36.11 | 502 | 76650 |
| **TOTAL** | **83.68** | **5940** | **172835** |

Since any map will include errors, it is critical to understand their nature because they can propagate to other data sets (*12, 13*). Bastin et al. (*1*) included an evaluation of the sampling and measurement errors attributable to estimation of forest area, but they did not do any attempt to validate the quality of the basal map used to define drylands (*2*). This could bring into question the validity of their results as a whole. A simple cross tabulation of a sample of predicted classes against their corresponding observed classes would have been enough (*14*). Many measures can be derived from such a confusion matrix and used to test the validity of the work done. Bastin et al. (*1*) should have tested this before designing their sampling scheme, and should have corrected it by excluding non-dry ecoregions and/or by re-computing the aridity index based on modern climatic data bases. In fact, other global assessments of the conservation status of dry forests (*15*) have taken these problems in consideration and have combined a corrected version of the aridity index (*6*) with the selection of ecoregions of interest (*3*).

Taken together all these limitations, it is likely that the 40 to 47% increase in the previous estimates of the extent of forest in drylands, and the potentially increase by 9% in the global area with over 10% tree canopy cover suggested by the results of Bastin et al. (*1*) will be substantially lower than claimed. This precludes the consideration and the use of the implications of their results to develop and reconsider strategies of mitigation and adaptation to global warming at a global scale.

## Acknowledgments

M.C., L.C. and A.E. are supported by project REMEDINAL3 [S2013/MAE-2719] from Comunidad de Madrid. A.E. is also supported by project ROOTS [CGL2015-66809-P] from the Spanish Ministry of Economy and Competitiveness. All the numbers and statistics cited in the main text have been computed using publicly available datasets and the open source programming language R. A script with all the computations is available upon request.

~~~
# **<u>“More than Drylands but Less of Dry forests”.</u>**
# Technical comment to be submitted to Science
# June, 07 2017.
# By Marcelino de la Cruz,
# Pedro F. Quintana-Acencio,
# Luis Cayuela
# Carlos I. Espinosa
# and Adrián Escudero
#
#########################################
#
#  R Script to document numbers, statistics and maps in the manuscript
#
##########################################
# 0) load the required R packages.
library(maptools)
library(rgdal)
library(raster)
library(rgeos)
#  --------------------------------------------------------------------------------
# 1) Read in and prerpare Bastin et al. 2017 data.
#  ----------------------------------------------------------------------------------
data_url<- “http://science.sciencemag.org/highwire/filestream/694256/
field_highwire_adjunct_files/1/aam6527_Bastin_Database-S1.csv.zip”
download.file(data_url, destfile=”aam6527_Bastin_Database-S1.csv.zip”)
unzip(“aam6527_Bastin_Database-S1.csv.zip”)
bastin <- read.csv(“aam6527_Bastin_Database-S1.csv”, header=T, sep=”;”)
# exclude 2 localities without coordinates (i.e., with NA ) bastin.naok<-
bastin[!is.na(bastin$location_x),]
# transform Bastin dataframe in an SpatialPointsDataFrame
coordinates(bastin.naok) = ~location_x+location_y
# Set projection attributes
pro <-”+proj=longlat +datum=WGS84 +no_defs +ellps=WGS84 +towgs84=0,0,0”
proj4string(bastin.naok)<-pro
# ----------------------------------------------------------------------------------------------------
# 2) Compute the aridity index (IA) based on recent climatic data
#     (Hijmans et al. 2015, Fick and Hijmans2017.) and check
#     how many plots of Bastin et al. are located in “Drylands”
# ----------------------------------------------------------------------------------------------------
# Download 2.5’ arc P data of worldclim 2.0
# The file is 68.5 MB so it takes some time to download
P_url<- “http://biogeo.ucdavis.edu/data/worldclim/v2.0/tif/base/wc2.0_2.5m_prec.zip”
download.file(P_url, destfile=”wc2.0_2.5m_prec.zip”)
unzip(“wc2.0_2.5m_prec.zip”)
# woldclim provides 12 layers of monthly Precipitation. Generate a stack object
# with all monthly layers
P_filenames<- paste(“wc2.0_2.5m_prec_”, c(paste(0,1:9, sep= “”), 11,12), “.tif”,sep= “”)
prec2.5<- stack(P_filenames)
# Compute annual precipitation as a raster layer
P2.5<- sum(prec2.5)
# Download 2.5’ arc PET data of Title and Bemmels (2017)
# The original data could be downloaded also from
# http://www.cgiar-csi.org/data/global-aridity-and-pet-database (Zomer et al. 2007, 2008)
# The file is 316.0 MB so it takes some time to download
pet_url<- “https://deepblue.lib.umich.edu/data/downloads/ms35t870p”
download.file(pet_url, destfile=”global_current_2.5arcmin_geotiff.zip”)
# Unzip PET data and generate raster object
unzip(“global_current_2.5arcmin_geotiff.zip”, files=”current_2-5arcmin_annualPET.tif”)
PET2.5<- raster(“current_2-5arcmin_annualPET.tif”)
# Compute the Aridity Index (AI = P/PET) as a raster layer
# The warning is because PET2.5 does not have data south of -60 ° (i.e. Antarctica, so, no problem)
PPET.ratio2.5<- P2.5/PET2.5
# Compute the map of dryland areas according to the criterion of UNCCD (i.e., IA <= 0.65)
drylands <- PPET.ratio2.5<0.65
# Calculate how many Bastin etal. plots are realy in areas with IA <0.65 (“1”) vs IA >=0.65 (“0”)
bastindry<-extract(drylands, bastin.naok)
table(bastindry)
#which proportions are these?
   table(bastindry)/sum(table(bastindry))
   #
# Visualize the distribution of “drylands” (IA >0.65) and the plots of Bastin et al.
# which are located in NO-drylands
plot(drylands)
points(bastin.naok[!as.logical(bastindry)&!is.na(bastindry),], col=2,pch=19, cex=0.2)
# Compute the average tree cover in plots located in “dry” (1) and in “non-dry” (2) plots
# and test for a difference among them
tapply(bastin.naok$tree_cover,list(factor(bastindry)), mean)
kruskal.test(bastin.naok$tree_cover,factor(bastindry))
# ------------------------------------------------------------------------------------------------------------------------------------
# 2.B) ALTERNATIVE: use Aridity Index from World atlas of desertification.
# and check how many plots of Bastin et al. are located in “Drylands”
# --------------------------------------------------------------------------------------------------------------------------------------
# Download 10’ arc aridity index map (Middleton & Thomas, 1997,
# World Atlas of desertification), provided by FAO
aridity_url<-
“http://www.fao.org/geonetwork/srv/en/resources.get?id=37040&fname=aridity.zip&access=private”
download.file(aridity_url, destfile=”aridity.zip”, mode=”wb”)
unzip(“aridity.zip”)
aridity<- raster(“./aridity/aridity”)
# Compute the map of dryland areas according to the criterion of UNCCD (i.e., IA <= 0.65)
drylands.FAO <-aridity<0.65
# How many plots are realy in areas with IA <= 0.65 (“1”) vs IA >0.65 (“0”)
bastindry.FAO<-extract(drylands.FAO, bastin.naok)
table(bastindry.FAO)
# wich proportions?
table(bastindry.FAO)/sum(table(bastindry.FAO))
# Visualize the distribution of “drylands” (IA >0.65) and the plots of Bastin et al.
# which are located in NO-drylands
plot(drylands.FAO)
points(bastin.naok[!as.logical(bastindry.FAO)&!is.na(bastindry.FAO),], col=2,pch=19, cex=0.2)
# ---------------------------------------------------------------------------------------------------------------------------------------------
# 3) Calculate overlap between Bastin et al. plots and Biomes distinguished by
#    WWF (Olson et al. 2001)
# ----------------------------------------------------------------------------------------------------------------------------------------------
# Download and pre-process Terrestrial Ecoregions (Olson et al. 2001) data from wwf
# # The file is 49.3 MB so it takes some time to download
wwf_url<-”<u>https://c402277.ssl.cf1.rackcdn.com/publications/15/files/original/official_teow.zip?1349272619</u>”
download.file(wwf_url, destfile=”official_teow.zip”, mode = “wb”)
unzip(“official_teow.zip”)
# Read wwf data into R as an Spatial PolygonsDataFrame object
ecoreg.wwf<- readOGR(“official”)
# transform the the variable “BIOME” in an informative factor. Levels are extrated from
# the metadata (wwf_terr_ecos.htm) included in the zip file
   ecoreg.wwf$BIOME<- factor(ecoreg.wwf$BIOME)
   biomeswwf<-c(“Tropical and Subtropical Moist Broadleaf Forests”,
              “Tropical and Subtropical Dry Broadleaf Forests”,
              “Tropical and Subtropical Coniferous Forests”,
              “Temperate Broadleaf and Mixed Forests”,
              “Temperate Conifer Forests”, “Boreal Forests/Taiga”,
              “Tropical and Subtropical Grasslands, Savannas and Shrublands”,
              “Temperate Grasslands, Savannas and Shrublands”,
              “Flooded Grasslands and Savannas”, “Montane Grasslands and Shrublands”,
              “Tundra”, “Mediterranean Forests, Woodlands and Scrub”,
                 “Deserts and Xeric Shrublands”, “Mangroves”, “class 98”,”class 99”)
levels(ecoreg.wwf$BIOME)<- biomeswwf
#How many plots are in each WWF biome
   #this could take some time
   biomes.wwf.bastin <- over(bastin.naok, ecoreg.wwf)
table(biomes.wwf.bastin$BIOME)
# Tabulate how many plots of Bastin et al. exists by wwf ecoregions and
# aridity zones (“0”: AI>=0.65, “1”: AI<0.65)
table.AIunep.wwf<-table(biomes.wwf.bastin$BIOME, bastindry)
   # Show first the non-dry biomes
orden<-c((1:16)[-c(2,7,8,10,12,13)],c(2,7,8,10,12,13))
table.AIunep.wwf[orden,]
# ------------------------------------------------------------------------------------------------------------------------------------------
# 3B) ALTERNATIVE: Calculate overlap between Bastin et al. plots and Biomes distinguished by
# TNC (a slightly modified version of WWF map)
# ------------------------------------------------------------------------------------------------------------------------------------------
   #Download and pre-process Terrestrial Ecoregions data from TNC (Olson and Dinerstein 2002)
   # # The file is 50.3 MB so it takes some time to download
   tnc_url<- “http://maps.tnc.org/files/shp/terr-ecoregions-TNC.zip”
   download.file(tnc_url, destfile=”terr-ecoregions-TNC.zip”, mode = “wb”)
   unzip(“terr-ecoregions-TNC.zip”, exdir =”./terr-ecoregions-TNC” )
   # Read TNC data into R as an Spatial PolygonsDataFrame object
   ecoreg.tnc<- readOGR(“./terr-ecoregions-TNC”)
   #How many plots are in each WWF biome
   #this could take some time
      biomes.tnc.bastin <- over(bastin.naok, ecoreg.tnc)
      table(biomes.tnc.bastin$WWF_MHTNAM)
   # Plots of Bastin et al by TNC ecoregions and aridity zones (“0”: AI>=0.65, “1”: AI<0.65)
      table.AIunep.tnc<-table(biomes.tnc.bastin$WWF_MHTNAM, bastindry)
      orden.tnc<- c(15,12,9, 10,1,3,16,5,4,8,   13, 14, 11, 7, 6, 2)
      table.AIunep.tnc[orden.tnc,]
# --------------------------------------------------------------------------------------------------------------------------------------------
# 4) OVERLAY between UNEP WCMC (Sorensen 2009) Drylands and
#    WWF (Olson et al. 2001) Terrestrial ecoregions
# ----------------------------------------------------------------------------------------------------------------------------------------------
# Download and pre-process UNEP WCMC (Sorensen 2009) Drylands data from UNEP
# There is not direct download of this data.
# You should visit this page:
# https://www.unep-wcmc.org/resources-and-data/world-dryland-areas-according-to-unccd-and-cbd-definitions
# click in “download data”, and after you fill in and submit a few details about yourself
#    and how you plan to use the data, and adhere to the terms and conditions,
#    a link to the dataset will be provided via email
# Place the downloaded zip file “Drylands_dataset_2007.zip” in your working directory
getwd ()
#    and continue
unzip(“Drylands_dataset_2007.zip”)
drylands.unep<- readOGR(“./Drylands_dataset_2007/Drylands_latest_July2014”)
   # 4 A) --------------------------------------------------------------------------------------------------------------
   # Show which portion of dry WWF biomes are ignored in Bastin et al, 2007 study:
   #    intersect each dry WWF biome with the dryland WCWP UNEP map. Exclude zones classified
   #    by Sorensen as “Presumed included: Drylands features but P/PET >= 0.65 “
   #    which have “NA in the field “HIX_DESC” and that have been ignored by Bastin et al. # This could take a few minutes
      # names of the dry biomes
   wwwf_dry_names <- biomeswwf[c(2,7,8,10,12,13)]
   dry.NOsampled.wwf <- list()
   for(name in wwwf_dry_names){
      dry.NOsampled.wwf[[name]] <- gDifference(ecoreg.wwf[ecoreg.wwf$BIOME==name,],
                                                               drylands.unep[!is.na(drylands.unep$HIX_DESC),])
   }
   # Area of dry WWF biomes ignored in Bastin et al:
   sapply(dry.NOsampled.wwf, area)
   # Proportion of the area of dry biomes ignored in Bastion et al.:
   sapply(dry.NOsampled.wwf, area)/aggregate(area(ecoreg.wwf[ecoreg.wwf$BIOME%in%wwwf_dry_names,]),
            by=list(ecoreg.wwf[ecoreg.wwf$BIOME%in%wwwf_dry_names,]$BIOME), FUN=sum)$x
   round(100*sapply(dry.NOsampled.wwf, area)/
            aggregate(area(ecoreg.wwf[ecoreg.wwf$BIOME%in%wwwf_dry_names,]),
            by=list(ecoreg.wwf[ecoreg.wwf$BIOME%in%wwwf_dry_names,]$BIOME), FUN=sum)$x,1)
   # --------------------------------------------------------------------------------------------------------------------------------------------------
   # Map of Dry forest in North and Central America
   #    Cyan: considered in UNEPP and Bastin et al.2017
   #    Yellow: Not considered in UNEPP and Bastin et al.2017
   data(wrld_simpl)
   dev.new(height=7, width=9)
   par(mar=rep(1,4))
      plot(wrld_simpl,ylim=c(-7,30), xlim=c(-120,-70), col=”grey”)
   plot(ecoreg.wwf[ecoreg.wwf$BIOME==”Tropical and Subtropical Dry Broadleaf Forests”,],
      col=”cyan”, add=T,border=”cyan”)
   plot(dry.NOsampled.wwf[[”Tropical and Subtropical Dry Broadleaf Forests”]],
      add=T,col=”yellow”, border=”yellow”)
   plot(wrld_simpl,add=T)
   box()
   axis(1, tck=0.02)
   axis(2, tck=0.02)
   axis(3, tck=0.02)
   axis(4, tck=0.02)
   text(x=rep(-120,4), y=c(30,20,10,0), labels=c(“30°”,”20°”,”10°”,”0°”))
   text (y=rep(-7, 6), x=c(-120,-110,-100,-90,-80,-70), labels=c(“120°”,”110°”,”100°”,”90°”,”80°”,”70°”))
   # ----------------------------------------------------------------------------------------------------------------------------------------
   # Map of Dry forest in the World
   #    Cyan: considered in UNEPP and Bastin et al.2017
   #    Yellow: Not considered in UNEPP and Bastin et al.2017
   dev.new(height=7, width=21)
   par(mar=rep(1,4))
   plot(wrld_simpl, col=”lightgrey”, xlim=c(-110, 130), ylim=c(-30, 30))
   plot(ecoreg.wwf[ecoreg.wwf$BIOME==”Tropical and Subtropical Dry Broadleaf Forests”,],
         col=”cyan”, add=T,border=”cyan”)
   plot(dry.NOsampled.wwf[[”Tropical and Subtropical Dry Broadleaf Forests”]],
         add=T,col=”yellow”, border=”yellow”)
   plot(wrld_simpl,add=T)
   box ()
   axis(1, tck=0.02)
   axis(2, tck=0.02)
   axis(3, tck=0.02)
   axis(4, tck=0.02)
   text(x=rep(-114,3), y=c(-20,0,20), labels=c(“20°”,”0°”,”20°”))
   text(y=rep(-36,5), x=c(-100,-50,0,50,100), labels=c(“100°”,”50°”,”0°”,”50°”,”100°”))
   # 4B) ----------------------------------------------------------------------------------------------------------------------------------
   #Show which portion of NO-dry WWF biomes are included in Bastin et al, 2007 study:
   #    intersect each NO-dry WWF biome with the dryland WCWP UNEP map
   # Exclude zones classified by Sorensen as “Presumed included: Drylands features but P/PET >= 0.65 “
   #    which have “NA in the field “HIX_DESC” and that have been ignored by Bastin et al.
   #    This could take a few minutes
   # Names of NO-Dry biomes
wwwf_NOdry_names <- biomeswwf[!biomeswwf%in%wwwf_dry_names]
NOdry.sampled.wwf2 <- list()
for(name in wwwf_NOdry_names){
   NOdry.sampled.wwf2[[name]] <- gIntersection(ecoreg.wwf[ecoreg.wwf$BIOME==name,],
                                                   drylands.unep[!is.na(drylands.unep$HIX_DESC),],
                                                   drop_lower_td=TRUE)
   }
   # Area of NO-dry WWF biomes included Bastin et al:
    sapply(NOdry.sampled.wwf2, area)
   # Proportion of the area of NO-dry biomes included in Bastion et al.:
   round(100*sapply(NOdry.sampled.wwf2, area)/
            aggregate(area(ecoreg.wwf[ecoreg.wwf$BIOME%in%wwwf_NOdry_names,]),\
            by=list(ecoreg.wwf[ecoreg.wwf$BIOME%in%wwwf_NOdry_names,]$BIOME), FUN=sum)$x,2)
# ---------------------------------------------------------------------------------------------------------------------------------------------------------
#    References
# ----------------------------------------------------------------------------------------------------------------------------------------------------------
# Bastin et al. 2017. The extent of forest in dryland biomes. Science 356, 635–638.
# Hijmans et al. 2005. Very high resolution interpolated climate surfaces for global
#    land areas. Int. J. Climatol. 25: 1965-1978.
# Fick, and Hijmans. 2017. WorldClim 2: new 1-km spatial resolution climate surfaces
#    for global land areas. Int. J. Climatol. 10.1002/joc.5086
# Middleton and Thomas, 1997. World atlas of desertification. United Nations
#    Environment Programme/Edward Arnold, London, 2nd edition.
# Olson et al. 2001 Terrestrial ecoregions of the world: a new map of life on Earth.
#    Bioscience 51,933-938.
# Olson and Dinerstein. 2002. The Global 200: Priority ecoregions for global
#    conservation. Annals of the Missouri Botanical Garden 89:125-126.
# Title and Bemmels. 2017. ENVIREM: An expanded set of bioclimatic and topographic
#    variables increases flexibility and improves performance of ecological niche
#    modeling. Ecography 10.1111/ecog.02880
# Zomer RJ, Trabucco A, Bossio DA, van Straaten O, Verchot LV, 2008. Climate Change
#    Mitigation: A Spatial Analysis of Global Land Suitability for Clean Development
#    Mechanism Afforestation and Reforestation. Agric. Ecosystems and Envir.
#    126: 67-80.
#
# Zomer RJ, Bossio DA, Trabucco A, Yuanjie L, Gupta DC & Singh VP, 2007. Trees and
#    Water: Smallholder Agroforestry on Irrigated Lands in Northern India. Colombo,
#    Sri Lanka: International Water Management Institute. pp 45. (IWMI Research
#    Report 122).
~~~

